# Biotic niche expansion constrains the fundamental abiotic niche: Evidence from experimental evolution

**DOI:** 10.64898/2026.04.27.720987

**Authors:** Gulsamal Askarova, Anna Skoracka, Ewa Puchalska, Mariusz Lewandowski, Jason Sexton, Lechosław Kuczyński

## Abstract

In an ever-changing world, organisms are subject to selective pressures that shape their ecological niches. Niche theory predicts that environmental heterogeneity selects for niche expansion, yet niches are inherently multidimensional, and expansion in one dimension may impose severe constrains on others. In this study, we employed rigorous experimental evolution to investigate these cross-dimensional trade-offs in the wheat curl mite, *Aceria tosichella*. By adapting replicated lineages to either stable (single-host) or alternating (two-host) environments for hundreds of generations, we successfully expanded the mites’ fundamental biotic host niche, enabling lineages to exploit diverse host species, including those unencountered during their evolutionary history. Crucially, however, this biotic generalization incurred a significant cost in the abiotic niche dimension. Lineages adapted to alternating hosts exhibited significantly reduced thermal tolerance compared to host specialists, which maintained superior performance across a wider thermal range. This trade-off appears to be driven by a combination of genetically based metabolic constraints and behavioral dispersal strategies. Our results provide compelling experimental evidence for the “Jack-of-all-trades is master of none” hypothesis across niche dimensions. We demonstrate that physiological trade-offs between biotic versatility and abiotic resilience strictly constrain the evolution of the multidimensional niche, with critical implications for forecasting species’ distributions and invasion potential under climate change.

## INTRODUCTION

In response to environmental selective pressures, the ecological niche, i.e., the set of conditions allowing for positive population growth, is expected to evolve. The niche may expand, contract, or shift depending on the nature of these environmental pressures. This results in organisms existing on a continuum from specialists, whose fitness is maximized in specific environments, to generalists, which maintain high fitness across diverse conditions (Levins, 1968; Sexton et al., 2017). However, the niche is inherently multidimensional, and organisms interact with their environment across multiple axes simultaneously, meaning that adaptation to one niche dimension may promote, constrain, or function independently of another (Carscadden et al., 2020; Hutchinson, 1957). Empirical evidence reflects this complexity, with scenarios ranging from the synergistic expansion of resource and thermal niches (Leonard & Lancaster, 2022) to severe trade-offs that compel specialization along a single axis (Litsios et al., 2014). Resolving whether and why broadening the niche in one dimension leads to functional trade-offs or rather promotes overall generalism is critical for forecasting the persistence, distribution, and invasion potential of populations in a changing world.

Despite the apparent advantages of ecological versatility, generalists rarely displace specialists in stable environments, implying that broad niches incur significant fitness costs (Bono et al., 2020). The “Jack-of-all-trades is master of none” hypothesis attributes these costs to antagonistic pleiotropy, where alleles beneficial in one environment are deleterious in others. Consequently, generalists are expected to be outcompeted by specialists within the latter’s optimal environment (Futuyma & Moreno, 1988). While some studies confirm these performance trade-offs (e.g., Cooper & Lenski, 2000; MacLean et al., 2004), others report neutral or even positive fitness correlations across environments instead (e.g., Draghi, 2021; Gompert et al., 2015; Magalhães et al., 2009, 2014). Alternative models propose that even in the absence of direct trade-offs, specialization may be favored by the efficiency of selection. Specialists experiencing consistent environmental pressures can fix beneficial alleles more rapidly than generalists (Hardy et al., 2020; Whitlock, 1996). Conversely, generalists may suffer from relaxed or conflicting selection pressures in any given environment, leading to the accumulation of deleterious mutations and reduced evolvability (Bono et al., 2020). Thus, generalism may be constrained not only by physiological costs but also by a slower rate of adaptation and an increased genetic load.

The mechanisms underlying niche expansion remain poorly understood, primarily due to the profound methodological challenges of studying the ecological niche in its multidimensional complexity. It is critical to account for both biotic and abiotic dimensions simultaneously, rather than seeking compromises strictly within a single environmental axis. This is particularly important because biotic interactions, such as host use, and abiotic, such as thermal limits, often demand fundamentally distinct, and potentially antagonistic, physiological mechanisms - as demonstrated by some comparative approaches (e.g. Bebber & Chaloner, 2022). Consequently, categorizing organisms strictly as overall ‘generalists’ or ‘specialists’ without actively decoupling these dimensions remains reductive. We argue that controlled empirical studies utilizing experimental evolution are essential to definitively disentangle causal, physiological trade-offs from independent evolutionary trajectories.

Our goal was to resolve whether niche expansion in one dimension promotes multidimensional generalism or whether functional constraints compel severe trade-offs. To achieve this, we experimentally broadened the fundamental biotic host niche of a phytophagous arthropod and examined the effect of this expansion on its performance under diverse host plant species and thermal conditions. Specifically, we subjected replicated populations of the wheat curl mite (*Aceria tosichella*) to rigorous experimental evolution over hundreds of generations in either a stable single-host environment or an alternating two-host environment.

We hypothesized that alternating biotic selection would promote host-generalism, thereby enhancing performance on novel host species. The central question of this study, however, is whether this biotic generalization constrains or facilitates abiotic tolerance, which we assessed by quantifying population growth across a broad temperature gradient. We prioritized this design in order to produce the high degree of biotic generalism required for such a test. We explicitly tested whether biotic versatility acts synergistically with thermal tolerance or incurs a physiological cost consistent with the “Jack-of-all-trades is a master of none” hypothesis.

Ultimately, we provide the first evidence from replicated, long-term experimental evolution, demonstrating that expanding the biotic niche reduced the fundamental abiotic resilience. By exposing these costs, we reveal that the evolution of multidimensional generalism can be strictly bound by severe physiological trade-offs.

## MATERIAL AND METHODS

### Study system

We utilized the wheat curl mite (*Aceria tosichella*; hereafter WCM) as our model system. WCM is an obligate phytophagous arthropod and a globally devastating agricultural pest, primarily due to its role as a vector for severe cereal viruses. While WCM comprises a cryptic species complex of genotypes with varying invasive capacities, we restricted our study to the MT-1 genotype, which is widely distributed and strongly associated with cultivated cereal fields (Skoracka et al., 2018). Crucially, its obligate dependence on living host tissue and rapid generation time make WCM an exceptionally tractable system for experimental evolution, allowing us to strictly enforce biotic niche expansion while subsequently isolating and testing abiotic (thermal) physiological constraints. Additional details regarding the biology, host range, and global distribution of WCM MT-1 are provided in Appendix S1.

### The concept of the study

*Aceria tosichella* and the general concept of the study is graphically presented in Figure 1.

**Figure 1.**
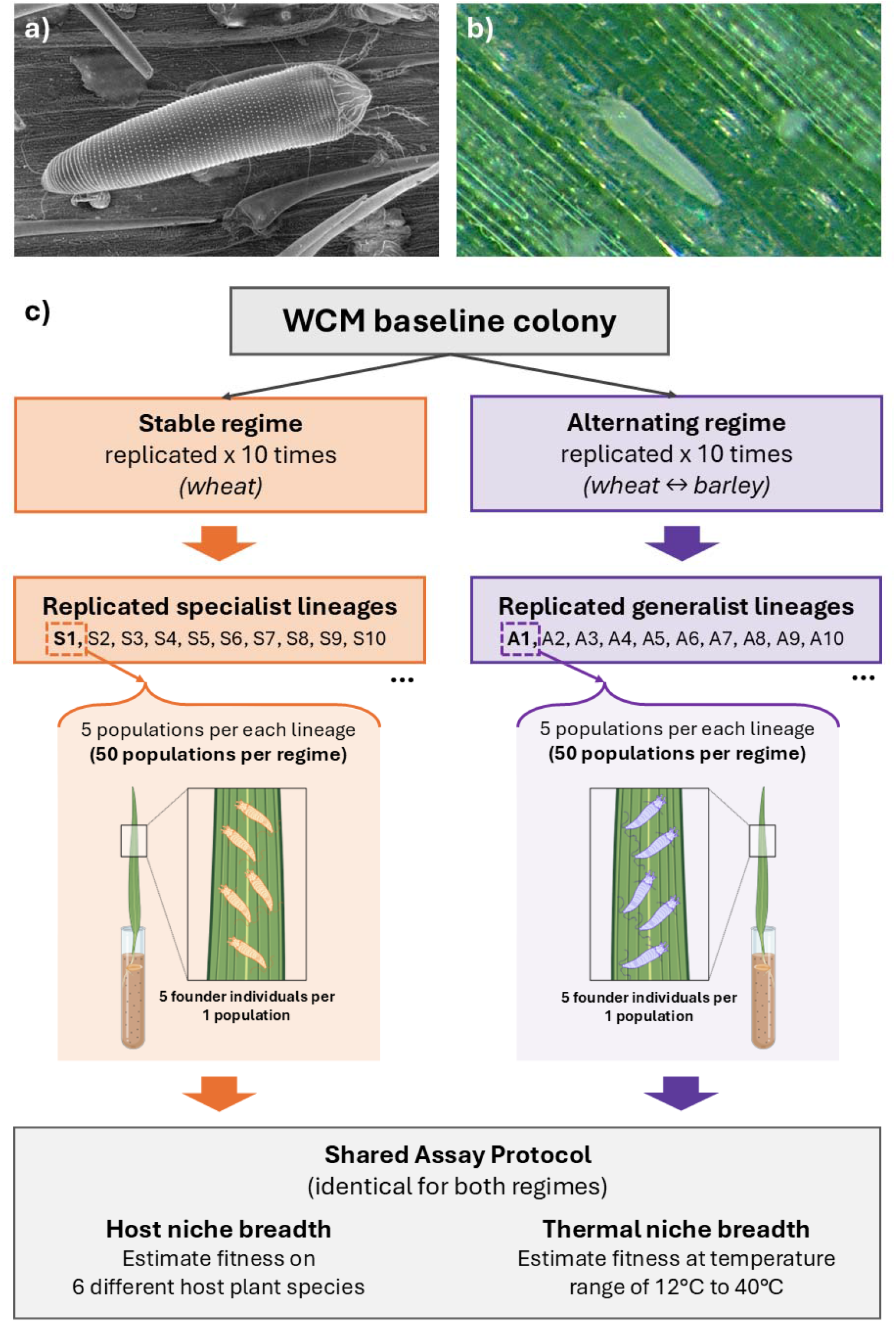
a) *Aceria tosichella* under a scanning electron microscope. b) *Aceria tosichella* under stereomicroscope. c) The concept of the study including experimental evolution in stable and alternating regimes, and testing host and thermal niche breadth of specialists and generalists.

#### The baseline colony

The baseline colony was established in November 2017 by mixing WCM MT-1 individuals collected from 24 field populations found in 10 geographically separated cereal growing areas in Poland, ensuring its genetic diversity. Before the build-up of the baseline colony, mite individuals from collected populations were barcoded using the mitochondrial cytochrome C oxidase subunit I (COI) sequence to confirm their MT-1 genotype. The baseline colony was maintained on wheat *Triticum aestivum* for five generations before it was used for experimental evolution (details about the creation of the baseline colony are provided in Skoracka et al. 2022).

#### Experimental evolution

We created two experimental regimes differing in the degree of stability of the biotic environment, and the host plant species acted as the biotic factor. We assumed that constant versus alternating biotic conditions should produce diverging adaptive responses (Kassen, 2002) resulting in a change in WCM host niche breadth. To do so, we subjected mites originating from the baseline colony to evolution in the following regimes:

i. stable conditions – on a single plant species: wheat (which served as a critical evolutionary control),
ii. alternating conditions – alternation of two plant species: wheat and barley *Hordeum vulgare*.

Evolution in each regime was replicated independently 10 times and lasted ca. 200 generations. In this way, from the same gene pool of sufficient genetic diversity we obtained 10 evolutionary lineages of ‘host specialists’ and 10 evolutionary lineages of ‘host generalists’.

Each lineage was established with approximately 300 WCM MT-1 individuals transferred from the baseline colony to clean potted plants and kept in separate isolators (i.e., contamination-protected cages). Each evolutionary regime was incubated separately in growth chambers under constant conditions of 24°C, 16:8 D/N, and 60% RH. Every 28 days (c.a. three generations on both wheat and barley), approximately 300 individuals from each replicated lineage were transferred to a pot with new clean plants according to the selection regime:

i. to wheat in stable environment,
ii. alternately to wheat or barley in alternating environment.

After the experimental evolution period lineages were maintained in a common garden environment on wheat for two full generations. Subsequently, we assessed the population growth rate in two separate experiments:

i. host niche breadth (biotic dimension): on a range of different plant species,
ii. thermal niche breadth (abiotic dimension): over a range of different temperatures.

### Estimating population growth rates

The experimental unit in our study was a population that developed on a single plant and was isolated from other populations. Each population was initiated with five females randomly selected and manually transferred from each replicated lineage (10 lineages evolved in stable and 10 lineages evolved in alternating regimes) to clean plants. We established five populations per each replicated lineage (Figure 1c). Because WCM reproduces parthenogenetically, both fertilized and unfertilized females can act as population founders (further details are provided in Appendix S1). To assess host niche breadth, we measured the growth rates of the populations on six host plant species, only two of which (wheat and barley) were previously involved in the experimental evolution. The other four were hosts that have never been encountered by experimentally evolving WCM: rye, *Secale cereale*; quackgrass, *Elymus repens*; tall-oatgrass, *Arrhenatherum elatius*; and smooth brome, *Bromus inermis*. All of these species are found within the potential range of the WCM. Rye is a different type of cereal, while the others grow at the edges of cultivated fields. To assess thermal niche breadth, we measured the growth rates of the populations on wheat over a temperature range of 12°C to 40°C, in four-degree increments. Immediately after the females were transferred to clean plants, populations were maintained in growth chambers at 24°C, 16:8 D/N, and 60% RH for the assessment of host niche breadth, and at the eight different temperatures, with the same photoperiod and humidity as above, for the assessment of thermal niche breadth. After 14 days in these conditions, the mites were counted.

## Statistical methods

All statistical analyses were performed in R version 4.4 (R Foundation for Statistical Computing, 2025) using the ‘mgcv’ to fit generalized additive mixed models (GAMM) and to run posterior sampling (Wood, 2017).

### Estimation of niche breadth

Generally, niche breadth is defined in terms of the range or diversity of conditions that allow the population to persist; i.e., ensure its non-negative instantaneous growth rates. To measure the host niche breadth, we used the normalized version of the Levins index (Levins, 1968), which in the context of this study measures the uniformity of the fitness distribution across *n* hosts:

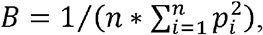

where *p*_*i*_ is a relative fitness on host *i*, such as 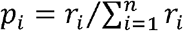, where *r*_*i*_ is the instantaneous population growth rate on the host *i*. When population growth rates are exactly the same for all hosts (and positive), the index takes its highest possible value (*B*=1). As the niche breadth decreases (i.e., the degree of host specialization increases), the index approaches zero. Populations with non-positive growth rates on all hosts were assigned an index of zero.

To measure the thermal niche breadth, we calculated the range of temperatures (*T*_*max*_ - *T*_*min*_) for which the instantaneous population growth rate (*r*) was positive.

### Statistical inference

To perform a statistical inference on niche breadth differences, we calculated posterior predictive distributions, which is a robust way to estimate confidence intervals for derived metrics (Gelman et al., 2004).

The procedure consisted of:

1. fitting GAMM models relating environmental conditions (host species or temperature) to population growth rates;
2. simulating a random draw from the posterior distribution of model parameters by using the Metropolis-Hastings sampler;
3. calculating predictions using the random draw;
4. estimating niche breadth by using these predictions;
5. estimating the effect size *Δ* by calculating the difference in niche breadth between experimental regimes (i.e. ‘Alternating’ vs. ‘Stable’);
6. repeating steps 2-5 10,000 times to obtain the distribution of sample statistics.

Then, for every considered statistic, its empirical 95% confidence intervals were calculated.

We used GAMM-s to assess relationships between environmental conditions (host or temperature) and population growth rates because of the method’s flexibility and built-in ability to perform posterior sampling. The model response was the finite population growth rate (λ), calculated for each population as a ratio of the number of mite individuals counted after 14 days of incubation to the number of females that were settled (which was always five). We specified a log link function in the GAMMs; consequently, the model’s linear predictor and parameter estimates represent the instantaneous population growth rate (*r*).

Growth rates in the two experimental regimes and on different host species were estimated using the following model:

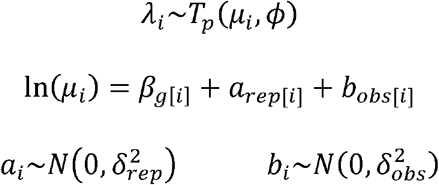

For each population *i*, its finite population growth rate *λ*_*i*_ was assumed to come from the compound Poisson–Gamma (Tweedie) distribution with the mean *μ*_*i*_, the variance power parameter *p* and dispersion parameter *ϕ*. The natural logarithm of the expected finite growth rate is the function of:

1. the group mean *β*_*g*[*i*]_, where double-indexing notation refers to the group identifier, which is a combination of host species and experimental regime (i.e., it is a 12-level factor: 6 hosts * 2 regimes);
2. the replicate effect, modelled as a normally distributed random intercept *a*_*rep*[*i*]_ with a mean at zero and variance 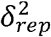, representing the variation arising from differences between independently evolved lineages for each experimental regime (20-level factor: 2 regimes * 10 replicates);
3. the observer effect, modelled as a normally distributed random intercept *b*_*obs*[*i*]_ with a mean at zero and variance 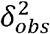, representing the observation error, i.e., the varying observer abilities to correctly estimate population size.

Growth rates in the two experimental regimes and at different temperatures were estimated using the following model:

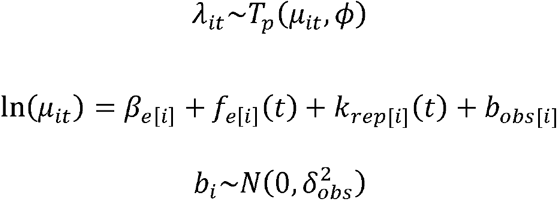

Now, growth rates have been estimated at a range of temperatures, so the finite population growth rate of the population *i* is also indexed by the temperature *t*. As in the previous model, it was assumed to come from the Tweedie distribution with the mean *µ*_*it*_, the variance power parameter *p* and dispersion parameter *ϕ*. The natural logarithm of the expected finite growth rate is the function of:

1. the group mean *β*_*e*[*i*]_, where e denotes an experimental regime (i.e., ‘Alternating’ or ‘Stable’);
2. the regime-specific smooth terms *f*_*e*[*i*]_ (*t*), which describes the general relationship between the temperature and population growth rate separately for each experimental regime (specified as a factor-smoother interaction);
3. the replicate-specific random smooths, modelled as a function *k*_*rep*[*i*]_ (*t*), representing the random deviation from the group-specific function estimated for each experimental regime (specified as a random factor-smoother interaction, using the tensor product for the continuous smoothers and random intercepts for the replicate-specific grouping parameters);
4. the observer effect modelled as a normally distributed random intercept *b*_*obs*[*i*]_ with a mean at zero and variance 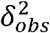, representing the observation error, i.e., the varying observer abilities to correctly estimate population size.

## RESULTS

Generally, the WCM populations responded to the level of biotic environmental variability. The stable host environment led to a narrow host range (i.e., host specialist) but a broad temperature tolerance. In contrast, the alternating host environment led to an expanded host range (i.e., host generalist) but a narrower thermal tolerance as compared to host specialists (Figures 2–3).

**Figure 2.**
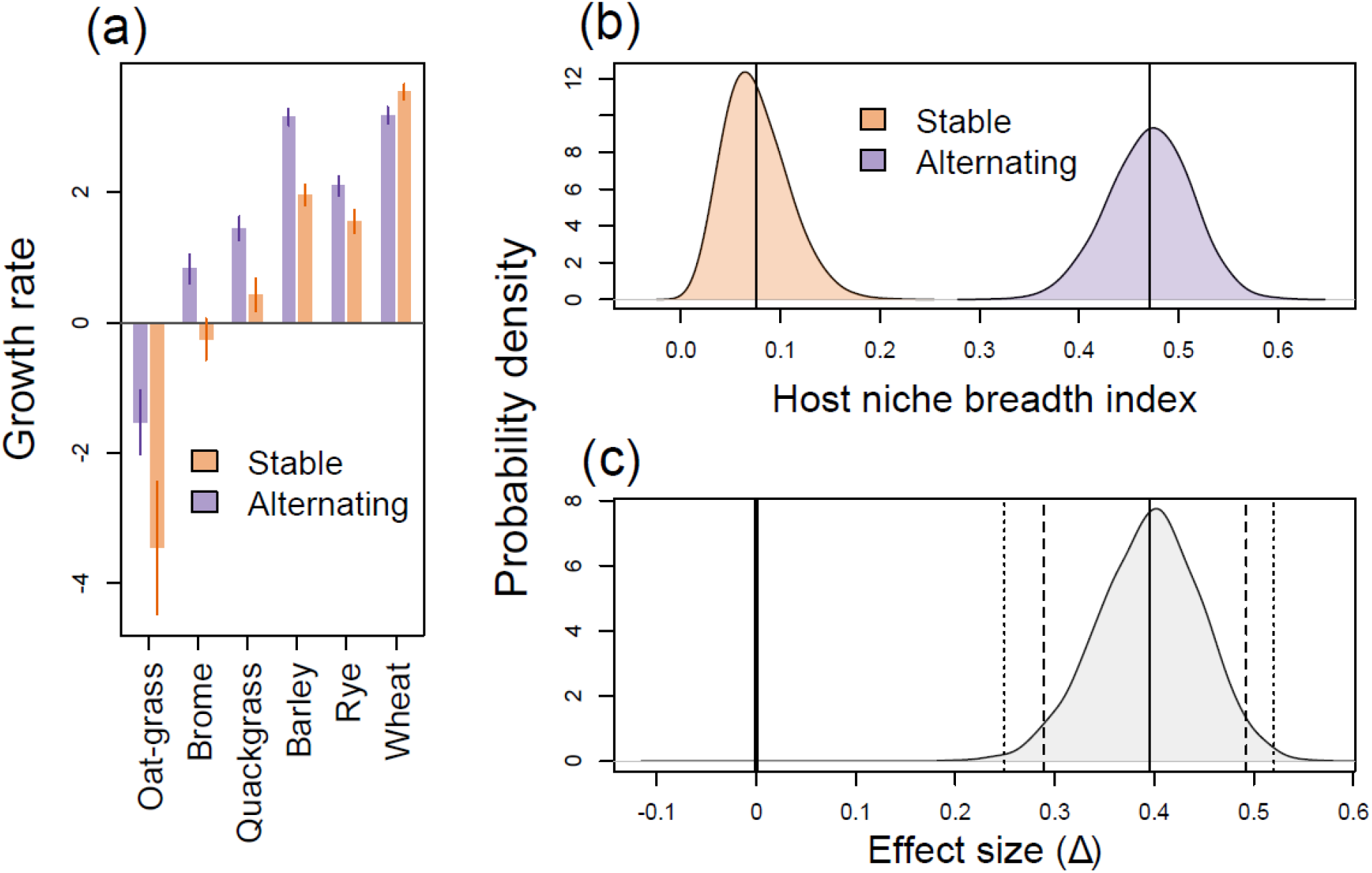
Host use patterns and estimated niche breadth for populations evolved in Stable and Alternating regimes. (a) Mean instantaneous population growth rates of experimental populations on the six plant species tested. Error bars represent 95% confidence intervals. (b) Posterior distributions of the host niche breadth index estimated for experimental populations evolved in Stable and Alternating environments. (c) Posterior distribution of the effect size, Δ, calculated as the difference between the host niche breadth estimated for experimental populations evolved in Alternating and Stable environments. Vertical lines denote 95% (dashed) and 99% (dotted) confidence intervals for the effect size. The solid line at zero indicates no effect.

**Figure 3.**
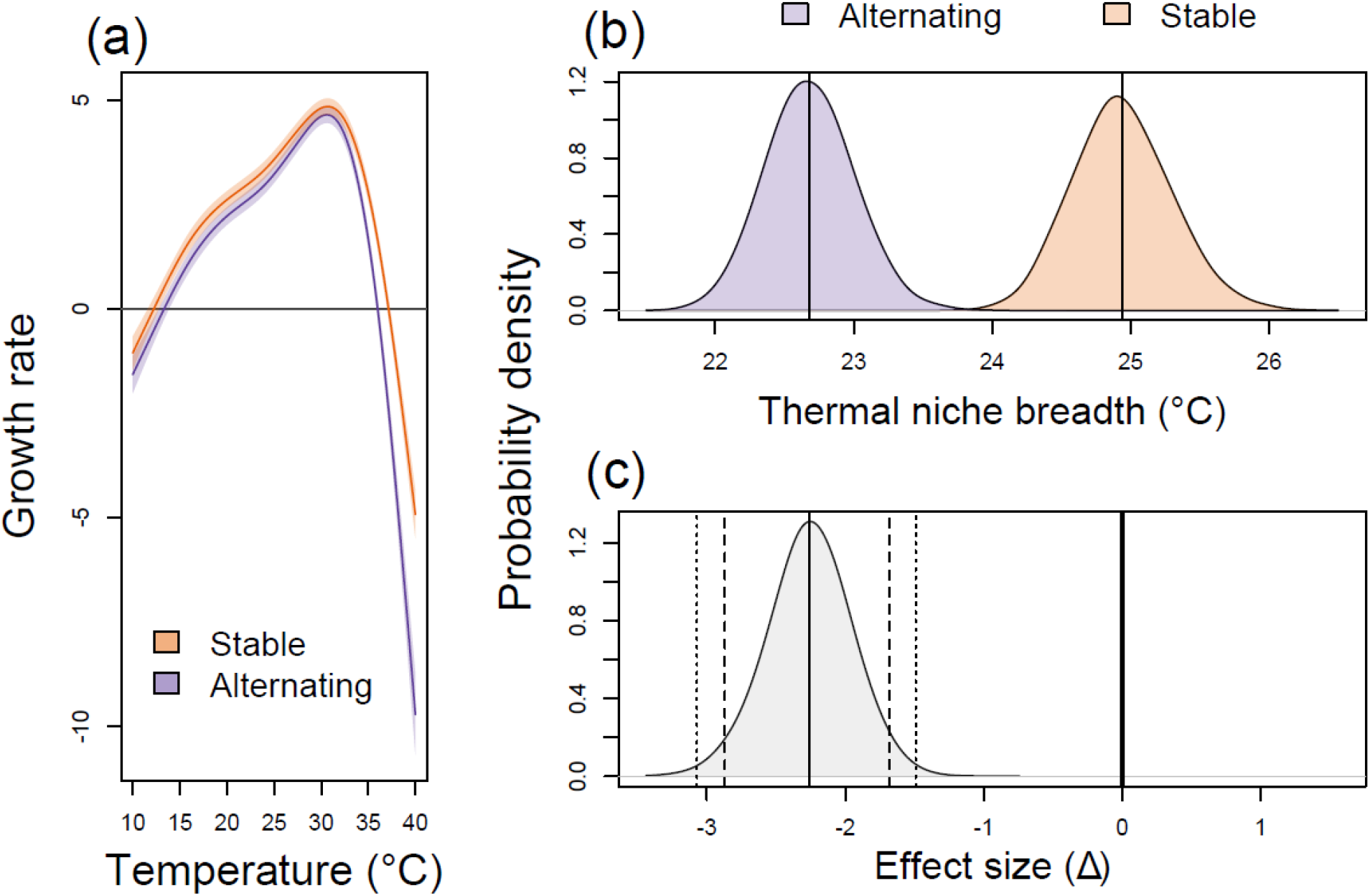
Thermal performance curves and estimated niche breadth for populations evolved in Stable and Alternating regimes. (a) Relationship between temperature and instantaneous growth rates of experimental populations that evolved in Alternating and Stable regimes. Lines represent the fitted GAMM functions, and shaded bands denote 95% confidence intervals. (b) Posterior distributions of the thermal niche breadth estimated for experimental populations evolved in Alternating and Stable regimes. (c) Posterior distribution of the effect size, Δ, calculated as the difference between the thermal niche breadth estimated for experimental populations evolved in Alternating and Stable environments. Vertical lines denote 95% (dashed) and 99% (dotted) confidence intervals for the effect size. The solid line at zero indicates no effect.

### Host niche breadth

Populations that evolved under alternating and stable host environments show different patterns of growth rates on the six plant species tested. In general, the populations that evolved in the alternating environment performed better than the populations that evolved in the stable host environment on all plant species except wheat (Figure 2a, Appendix S2: Table S1).

The Levins host niche breadth index was 0.47 (95% CI: 0.39-0.55) for the alternating regime and 0.08 (0.02-0.15) for the stable regime (Figure 2b). The effect size, i.e., the difference between alternating and stable regimes is 0.39 (0.29-0.49) and can be considered highly significant: the posterior distributions of the index values for both groups do not even overlap (Figure 2c).

### Thermal niche breadth

The fitted GAMM successfully described the relationship between temperature and population growth rates (Figure 3a, Appendix S2: Table S2, Figure S1). Populations that evolved under the alternating host regime have a narrower thermal niche compared to those that evolved under the stable host regime (22.7°C vs. 24.9°C, Table 1, Figure 3b). The effect size, i.e., the difference in thermal niche breadth, is −2.25°C and both its 95% and 99% confidence intervals does not contain zero (−2.86 to −1.67 and −3.06 to −1.48, respectively, Figure 3c).

**Table 1.** Thermal niche parameters for both experimental regimes. *T*_*min*_ and *T*_*max*_ determine the range of temperatures that allow for positive population growth; this range defines the thermal niche breadth. *T*_*opt*_ is the temperature at which the population growth reaches its maximum, and *λ*_*max*_ is the finite population growth rate at that temperature (i.e. the maximum expected growth rate). CI denotes 95% confidence intervals. Units are °C, except for the growth rate *λ*_*max*_, which is unitless.

| Parameter | Alternating regime |  | Stable regime |  |
| --- | --- | --- | --- | --- |
|  | Mean | 95% CI | Mean | 95% CI |
| $T_{min}$ | 13.3 | 12.7 – 13.8 | 12.2 | 11.6 – 12.7 |
| $T_{max}$ | 36.0 | 35.8 – 36.1 | 37.1 | 37.0 – 37.3 |
| $T_{opt}$ | 30.6 | 30.4 – 30.8 | 30.7 | 30.5 – 30.9 |
| $\lambda_{max}$ | 105.3 | 86.3 – 127.9 | 128.4 | 105.6 – 155.6 |
| $T_{max} - T_{min}$<br>(niche breadth) | 22.7 | 22.1 – 23.3 | 24.9 | 24.3 – 25.6 |

## DISCUSSION

### Cross-dimensional trade-offs constrain niche expansion

Organisms experience selection pressures across multiple, often independent, ecological dimensions. The evolution of niche breadth therefore depends on whether adaptive gains along one axis incur costs along others. Our primary objective was to generate generalist phenotypes to explicitly test for cross-dimensional physiological trade-offs. By subjecting populations to alternating host environments, we successfully evolved an expansion of the biotic niche, enhancing the mites’ ability to exploit diverse host species, including those unencountered during their evolutionary history. Crucially, this biotic generalization incurred a severe abiotic penalty: host-generalists exhibited significantly reduced thermal tolerance compared to host-specialists. By becoming ‘Jacks’ of the biotic environment, the mites inherently lost their ‘Mastery’ of the abiotic environment, providing compelling experimental evidence that fundamental trade-offs can strictly constrain the simultaneous evolution of overall niche breadth.

This profound constraint likely emerges because selection in our system acted across functionally distinct ecological axes. While synergistic niche expansions have been documented - e.g., resource generalism increasing thermal tolerance via shared stress-response pathways (Leonard & Lancaster, 2022) - these positive correlations typically occur within similar niche types involving interacting abiotic climatic factors (e.g. Bonetti & Wiens, 2014; Hernandez et al., 2023; Liu et al., 2020a). In contrast, traversing the boundary between biotic interactions (host use) and abiotic limits (temperature) exposes deeper physiological conflicts (Bebber & Chaloner, 2022; Chaloner et al., 2020). Our results suggest that the evolutionary effort required to maintain high performance across multiple hosts comes at the cost of a reduced abiotic niche. Thus, the physiological architecture required for biotic versatility appears fundamentally incongruous with the maintenance of broad thermal resilience.

Although our experimental framework focuses on two specific niche dimensions, thermal and host-plant niches represent the primary, defining axes governing the performance and distribution of phytophagous arthropods. Temperature directly controls key life-history traits - such as development, reproduction, and survival - thereby setting broad physiological and geographic limits (Ma et al., 2021). Concurrently, host-plant associations determine nutrient acquisition and exposure to defensive compounds, shaping fitness and population dynamics at local scales (Forister et al., 2012, 2015). Crucially, our findings underscore that these fundamental axes are not independent. Just as temperature can alter host plant chemistry (Sun et al., 2023), our results demonstrate that adaptation to diverse hosts actively constrains thermal tolerance, highlighting the deeply interconnected nature of the multidimensional niche.

### Drivers of the generalist trade-off

The fitness trade-offs we observed likely have a genetic basis rooted in antagonistic pleiotropy, where alleles that enhance performance across diverse host defenses simultaneously reduce thermal resilience (Agrawal, 2019; Laughlin & Messier, 2015). Our recent genomic evidence in WCM supports this, revealing distinct molecular profiles between specialist and generalist lineages and suggesting that the transition to a generalist strategy requires significant genomic restructuring. Generalism in WCM relies on specialized systems to manage the physiological stress arising from toxic compounds present in suboptimal or refuge hosts (Laska et al., 2021; Laska-Modzelewska et al., 2026). In phytophagous arthropods, the regulatory pathways used to navigate plant defenses often overlap with those governing temperature response (Ali & Agrawal, 2012; García-Robledo & Horvitz, 2012). Host generalism requires costly, versatile detoxification systems (Berenbaum, 1995; Heidel-Fischer & Vogel, 2015), whereas thermal tolerance depends on energy-intensive mechanisms like heat shock proteins and membrane stabilization (Feder & Hofmann, 1999). Thus, our results suggest that metabolic investment required to maintain broad detoxification pathways directly limits the resources available for abiotic stress-tolerance.

Beyond physiology, behavioral adaptations may further explain why biotic generalists exhibited reduced abiotic tolerance. Our recent results indicate that generalist WCM genotypes disperse more frequently and show heightened sensitivity to host cues compared to specialists (Zalewska et al., 2026). This aligns with theoretical expectations that generalists rely on high mobility to exploit heterogeneous landscapes (Ravigné et al., 2024). This heightened mobility likely serves as a behavioral buffer, allowing generalists to spatially “escape” thermal extremes rather than enduring them physiologically. Consequently, the evolution of biotic generalism appears to facilitate a transition from physiological endurance to behavioral avoidance, fundamentally reshaping the organism’s multidimensional niche.

### Agricultural management as a driver of range dynamics

Host range and thermal tolerance jointly shape the distributions of phytophagous arthropods, yet the relationship between diet breadth, thermal limits, and geographic range size remains notoriously inconsistent. Contrary to expectations, empirical evidence suggests that broad host use does not reliably predict range size (Lancaster, 2020; Stewart et al., 2015). This is often because climatic extremes constrain persistence more strongly than host availability, particularly in temperate systems where seasonality dictates geographic limits (Peguero et al., 2017; Rasmann et al., 2014). These mixed results reflect a complex interplay of factors, including spatial variation in host specificity and abiotic requirements (Arnal et al., 2019; Scriber, 2010).

Our results provide a physiological and evolutionary explanation for these inconsistencies: we demonstrate that in WCM, an expanded biotic niche comes at the direct cost of reduced thermal tolerance. This trade-off suggests that host-generalist lineages, despite their ecological versatility, possess a reduced capacity to colonize thermally variable or extreme environments. Consequently, these biotically versatile phenotypes may be confined to narrower climatic zones. In contrast, host-specialists - by maintaining superior thermal breadth - may be better equipped to inhabit broader geographic ranges, provided their primary cereal hosts are available.

From an applied perspective, our results indicate that human-induced variation in agricultural environments may fundamentally shape pest evolution and invasion potential. Management practices such as monocultures may favor host specialization coupled with broad thermal tolerance. Conversely, crop rotation or post-harvest host loss creates fluctuating environments that select for host range expansion, but our work reveals that this expansion is not cost-free, and that biotic generalism comes at the expense of abiotic resilience. This trade-off suggests that agricultural heterogeneity does not simply foster generalism; it drives an evolutionary dichotomy that determines whether a pest is ultimately limited by its host plant or its climate. This mirrors “rotation resistance” observed in other major pests, such as the Western Corn Rootworm (Gray et al., 2009; Levine et al., 2002), where agricultural heterogeneity drives rapid, and often costly, niche shifts.

Ultimately, because invaders tend to conserve their climatic niches in new regions (Liu et al., 2020b), broad host generalism does not inherently guarantee invasion success. Instead, our findings highlight how climatic tolerance can override the expected advantages of dietary breadth. If the evolution of biotic versatility inherently compromises thermal resilience, then the lineages most capable of switching to novel crops may simultaneously be the most vulnerable to the increasing frequency of thermal extremes. Conversely, the evolution of host specialization promoted by stable monocultures retains superior thermal resilience. As a consequence, host specialists may exhibit enhanced expansion potential under shifting global temperature regimes. Recognizing these cross-dimensional trade-offs is therefore critical for forecasting the spatial dynamics of agricultural pests under the dual pressures of intensified land use and a changing climate.

### Synthesis and future perspectives

Our results demonstrate that the evolution of biotic generalism incurs a significant abiotic cost: the capacity to exploit diverse hosts trades off against the ability to tolerate thermal variability. This underscores the multidimensional nature of the ecological niche, where resilience on one axis does not predict robustness on another. Consequently, accurately forecasting species distributions, invasion potential, community assembly, and climate-change vulnerability requires ecological models that explicitly distinguish between these niche dimensions. We suggest that future ecological trajectories will be shaped by this dichotomy: host generalists may readily establish in novel communities but require stable climates to persist, whereas specialists may possess the physiological robustness to endure climatic shifts but remain constrained by host availability. To fully map this multidimensional landscape, future research should employ reciprocal experimental evolution - subjecting populations to fluctuating thermal regimes - to test whether abiotic variability conversely selects for generalized host tolerance. Disentangling the underlying genetic architecture will be critical to determine whether abiotic adaptation facilitates host range expansion or if the trade-offs we observed are strictly bidirectional. Ultimately, biological responses to a rapidly changing world will strongly depend on this critical interplay between a species’ host breadth and its thermal limits.

## Supporting information

Supplementary Results

## ACKNOWLEDGMENTS

We are grateful to Jarosław Raubic, Agata Magdańska, Krystian Zadrowski, Maja Malinowska, Radosław Gmyrek, Wiktoria Szydło, and Katarzyna Kaszewska for their assistance with laboratory experiments. We thank the company DANKO Hodowla Roślin Sp. z o.o. for the seeds of *Triticum aestivum, Hordeum vulgare*, and *Secale cereale*. This study was supported by the National Science Centre (NSC), Poland, research grant no. 2021/41/B/NZ8/01703.

## Author Contributions

LK and AS conceptualized and designed the study with GA, EP, ML assistance; GA, EP, ML, AS ran experiments and collected the data; LK performed statistical analyses; GA, LK, AS, JS critically interpreted the results; AS, GA, JS, LK wrote the manuscript with assistance of EP, ML; All authors contributed substantially to revisions, read and approved the final version of the manuscript.

## Data accessibility statement

The data that support the findings and the code used to produce the results of this study are openly available on GitHub (https://github.com/popecol/niche_breadth) and Zenodo (https://doi.org/10.5281/zenodo.17910844).

