## Supplementary Methods for "Biotic niche expansion constrains the fundamental abiotic niche: Evidence from experimental evolution"

**APPENDIX S1**

**Biotic niche expansion constrains the fundamental abiotic niche: Evidence from experimental evolution**

Gulsamal Askarova, Anna Skoracka*, Ewa Puchalska, Mariusz Lewandowski, Jason Sexton, Lechosław Kuczyński

*Ecology*

* corresponding author

**Supplementary Methods**

**The WCM MT-1 Genotype Study System**

The obligate phytophagous mite *Aceria tosichella* (wheat curl mite, hereafter WCM) genotype MT-1 is strongly associated with cultivated cereal fields. Its primary recorded host plants include wheat, barley, rye, and triticale (Skoracka et al., 2018, 2022). However, this genotype can persist for several generations on alternative grass species, such as brome or quackgrass, which function as temporary stepping stones when the primary hosts senesce or become otherwise unavailable (Laska et al., 2021; Skoracka et al., 2018). The MT-1 lineage is globally distributed, having been documented across the Nearctic, Palearctic, and Australasian regions (Skoracka et al., 2014). The mite's microscopic size allows it to infest agricultural commodities undetected, while its capacity for long-distance aerial dispersal via wind currents and its broad thermal tolerance significantly enhance its invasive potential (Kuczyński et al., 2016; Navia et al., 2013).

WCM serves as an exceptionally tractable study system for experimental evolution due to its ease of laboratory maintenance, rapid population growth, and short generation time (e.g., egg-to-egg developmental time at 24°C is approximately 8.7 days) (Karpicka-Ignatowska et al., 2019, 2021; Skoracka et al., 2022). The life cycle of WCM comprises four distinct stages: egg, larva, nymph, and adult. Each active immature stage (larva and nymph) is followed by a period of quiescence, i.e., state of immobility lasting from several hours to a few days, prior to molting into the subsequent stage. Males actively patrol these quiescent stages, identifying and guarding nymphs that will molt into mature females. Around these female nymphs, males deposit spermatophores, which the females subsequently collect upon emerging as adults. Adult females are readily distinguished from other life stages and from males by their larger size and characteristic body shape. Furthermore, WCM reproduces via arrhenotokous parthenogenesis (Miller et al., 2012), wherein haploid males develop from unfertilized eggs and diploid females develop from fertilized eggs. This reproductive system allows a single unfertilized female — dispersed, for instance, via wind currents as a nymph — to found a novel population. By producing haploid sons and subsequently collecting their spermatophores, the foundress can then produce diploid female progeny. This strategy facilitates exceptionally rapid colonization and demographic expansion when the mite encounters isolated hosts or novel environments.
