## Supplementary Results for "Biotic niche expansion constrains the fundamental abiotic niche: Evidence from experimental evolution"

**APPENDIX S2**

**Biotic niche expansion constrains the fundamental abiotic niche: Evidence from experimental evolution**

Gulsamal Askarova, Anna Skoracka*, Ewa Puchalska, Mariusz Lewandowski, Jason Sexton, Lechosław Kuczyński

*Ecology*

* corresponding author

**Supplementary Results**

**Table S1.** Results of the GAMM estimating population growth rates across host plants and experimental regimes. Parametric 'Estimates' represent the mean instantaneous population growth rate for each host-regime combination. Random effect 'Estimates' represent the standard deviation of the normal distribution of random intercepts for replicates and observers. The model explains 88.8% of the deviance.

| **Parametric coefficients for hosts in the Alternating regime** | | | | |
| --- | --- | --- | --- | --- |
|  | Estimate | SE | t | p |
| Wheat | 3.18 | 0.063 | 50.1 | <0.0001 |
| Barley | 3.15 | 0.064 | 49.3 | <0.0001 |
| Rye | 2.10 | 0.079 | 26.6 | <0.0001 |
| Quackgrass | 1.44 | 0.093 | 15.5 | <0.0001 |
| Brome | 0.82 | 0.112 | 7.4 | <0.0001 |
| Oatgrass | -1.53 | 0.252 | -6.1 | <0.0001 |
| **Parametric coefficients for hosts in the Stable regime** | | | | |
|  | Estimate | SE | t | p |
| Wheat | 3.54 | 0.061 | 58.1 | <0.0001 |
| Barley | 1.96 | 0.082 | 23.9 | <0.0001 |
| Rye | 1.55 | 0.091 | 17.0 | <0.0001 |
| Quackgrass | 0.43 | 0.127 | 3.4 | 0.0007 |
| Brome | -0.26 | 0.160 | -1.6 | 0.1078 |
| Oatgrass | -3.46 | 0.521 | -6.6 | <0.0001 |
| **Random factors** | | | | |
|  | Estimate | edf | F | p |
| Replicate | 0.14 | 13.2 | 2.66 | <0.0001 |
| Observer | 0.05 | 2.2 | 1.18 | 0.0598 |

**Table S2.** Parameters of the GAMM modeling the relationship between temperature and population growth rate in two experimental regimes. Parametric coefficients represent the regime-specific intercepts (mean instantaneous growth rates). Smooth terms for regimes estimate the non‑linear temperature response curves. Smooth terms for replicates represent the random variation in the temperature response shape among replicates (factor-smooth interactions). For the random factor (Observer), 'Estimate' is the standard deviation of the random intercepts, while the test statistics (edf, F, p) refer to the penalized smooth term significance. The model explains 95.4% of the deviance.

| **Parametric coefficients for experimental regimes** | | | | |
| --- | --- | --- | --- | --- |
|  | Estimate | SE | t | p |
| Alternating | 0.57 | 0.117 | 4.8 | <0.0001 |
| Stable | 1.66 | 0.103 | 16.2 | <0.0001 |
| **Smooth terms for experimental regimes** | | | | |
|  |  | edf | F | p |
| Alternating |  | 5.0 | 780.6 | <0.0001 |
| Stable |  | 5.0 | 830.5 | <0.0001 |
| **Smooth terms for replicates** | | | | |
|  |  | edf | F | p |
| Replicate |  | 10.0 | 0.13 | 0.0561 |
| **Random factor** | | | | |
|  | Estimate | edf | F | p |
| Observer | 0.21 | 3.85 | 15.4 | <0.0001 |

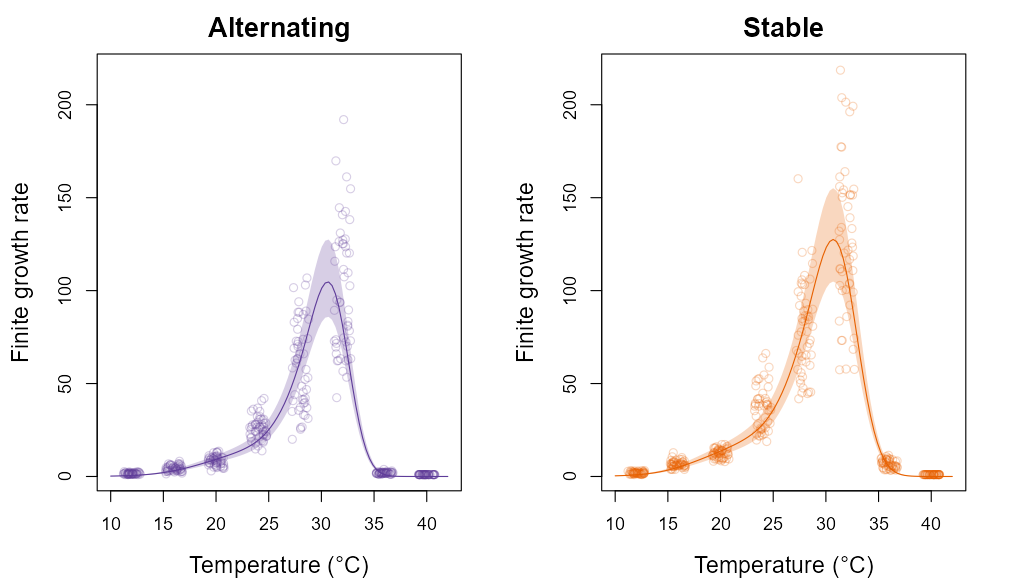

**Figure S1.** Relationship between temperature and finite population growth rates of experimental populations that evolved in alternating and stable environments. Points are observed values; lines are fitted functions and shaded bands are 95% confidence intervals.
